# *Contact*Blot: Microfluidic Control and Measurement of Cell-Cell Contact State to Assess Contact-Inhibited ERK Signaling

**DOI:** 10.1101/2023.11.06.565857

**Authors:** Yizhe Zhang, Isao Naguro, Hiroki Ryuno, Amy E. Herr

## Abstract

Extracellular signal-regulated kinase (ERK) signaling is essential to regulated cell behaviors, including cell proliferation, differentiation, and apoptosis. The influence of cell-cell contacts on ERK signaling is central to epithelial cells, yet few studies have sought to understand the same in cancer cells, particularly with single-cell resolution. To acquire same-cell measurements of both phenotypic (cell-contact state) and targeted-protein profile (ERK phosphorylation), we prepend high-content, whole-cell imaging prior to endpoint cellular-resolution western blot analyses for each of hundreds of individual HeLa cancer cells cultured on that same chip, which we call *contact*Blot. By indexing the phosphorylation level of ERK in each cell or cell-cluster to the imaged cell-contact state, we compare ERK signaling between isolated and in-contact cells. We observe attenuated (∼2×) ERK signaling in HeLa cells which are in-contact *versus* isolated. Attenuation is sustained when the HeLa cells are challenged with hyperosmotic stress. Our findings show the impact of cell-cell contacts on ERK activation with isolated and in-contact cells, while introducing a multi omics tool for control and scrutiny of cell-cell interactions.

## INTRODUCTION

Cell behaviors are constantly affected by signals from the outside microenvironment such as neighbor cells and extracellular matrix components. As a fundamental cell-cell interaction, physical contact between epithelial cells has been reported to inhibit a wide variety of critical cell activities in culture, including cell growth, proliferation, autophagy, phagocytosis, movement, adhesiveness, and plasticity of stem cells.^[1—8]^ While knowledge is accumulating towards a comprehensive understanding of contact-inhibited cell processes, research findings have suggested the involvement of pathways of mitogen-activated protein kinases (MAPKs) in the molecular mechanisms of contact inhibition.^[9—11]^ MAPKs (e.g., ERK1/2, p38α/β/γ/δ, JNK1/2/3) underpin myriad cell activities. Through rapid phosphorylation, MAPKs regulate a wide range of cell processes, including proliferation, differentiation, stress responses, apoptosis, and immune response.^[12—18]^ Hence, comparative analysis of MAPK activation in isolated *versus* in-contact paired cells should provide additional insight into signal regulation relevant in physiology and pathology.

Studies suggest that contact-inhibition is a local phenomenon rather than a global effect across a layer of cells,^[9]^ thus creating interest in scrutinizing MAPK activation for a variety of cell-contact state configurations with single-cell resolution. However, the majority of contact-inhibition studies have been performed in bulk. Also in bulk, behaviors of high-density *versus* low-density cells or interior *versus* peripheral cells in a monolayer have been compared,^[1, 9, 19]^ with finer resolution analyses stymied by technological limitations in single-cell culture and analyses. A key challenge is that adherent cells tend to grow in clusters in bulk, even at a low seeding density.^[20, 21]^ Depending on the cell-spreading behavior, mixed cell-contact states can appear within each cell-contact group. Hence, it is challenging to obtain information from unambiguously isolated *versus* unambiguously in-contact cells. On the other hand, the lack of effective techniques to detect MAPK activation in mammalian cells with single-cell resolution limits the understanding in this field: conventional assessment of phosphorylation through mass spectrometry or western blotting provides ensemble measurement from a population of cells.^[22, 23]^ Flow cytometry can interrogate millions of single cells for MAPK phosphorylation with high sensitivity. Yet the need for single-cell suspensions leads to the loss of cell-contact information. And the fixative used in the sample preparation can interfere with target epitopes.^[24]^ Newer single-cell resolution assays based on immunocytochemistry or mass spectrometry of phosphorylation offer limited throughput and require intensive data analysis.^[25-29]^

To obtain precision measurement of MAPK in isolated *versus* in-contact cells, we introduce a multi modal microfluidic assay, called *contact*Blot for brevity: the first mode is whole-cell imaging after on-chip cell culture in a microwell followed by a second mode which is an end-point single-cell or contact-cell western blot (μWB), wherein the microwell provides short-term cell-culture conditions and isolates the cell or cell pair.^[30, 31]^ Here we utilize bright-field whole-cell imaging to visually determine isolated *versus* in-contact status for each cell. We applied the *contact*Blot to HeLa cells to assess ERK phosphorylation under different osmotic conditions (a widely used stress-stimulus for MAPK-signaling studies), and observed differential phosphorylation levels of ERK in isolated *versus* in-contact cells under each osmotic condition. The in-contact cells exhibit a lower level of ERK activation compared to the isolated cells.

## EXPERIMENTAL SECTION

### Chemicals and Materials

N-[3-[(3-benzoylphenyl)formamido]propyl]methacrylamide (BPMAC) was purchased from PharmAgra Labs (Brevard, NC, USA). Primary antibodies for β-tubulin (rabbit; ab6046) were purchased from Abcam (Cambridge, UK). Primary antibodies for phosphorylated extracellular signal-regulated kinase (p-ERK) (rabbit; 4370S) and for phosphorylated p38 MAPK (p-p38) (rabbit; 4511) were purchased from Cell Signaling Technology (Danvers, MA, USA). Anti-rabbit secondary antibodies (Donkey, AlexaFluor 555; A31572) were purchased from Life Technologies (Carlsbad, CA, USA). Mylar masks with microwell features were purchased from CAD/Art Services (Bandon, OR, USA).

### Design and Fabrication of *contact*Blot Device

Fibronectin (FN) is used as the extracellular matrix (ECM) protein to functionalize microwells on the *contact*Blot device for on-chip cell culture.^[30]^ Other ECM proteins (for example, collagen, gelatin, laminin) can also be used based on the same principle. FN-functionalization of PA gels is completed at the gel polymerization step. Specifically, FN solution of 10 μg mL^-1^ is added to the precursor solution (mainly composed of acrylamide, bis-acrylamide, benzophenol-methacrylamide, APS, and TEMED) of the PA gel. The FN and PA gel precursor solution has a viscosity similar to water and is uniformly applied onto the SU-8 mold. A glass microscope slide is treated with silane to enhance surface hydrophilicity layered on top of the SU-8 mold thus forming what will be the foundational substrate of the *contact*Blot device. The SU-8-precursor-glass slide sandwich is kept at room temperature for ∼1 h to allow for polymerization. Once the polymerization is completed, the gel is peeled off from the SU-8 mold leaving an open microfluidic device consisting of an open microwell array stippled into a mini-PA-gel-slab on the microscope slide. Optionally, UV photocrosslinking can be implemented before or after the mold-release step to form covalent bonds between FN and the gel. To facilitate gel detachment from the SU-8 mold, the SU-8-gel-glass slide sandwich can be immersed in deionized water for ∼5 min prior to peeling.

### Cell Culture on the *contact*Blot Device

HeLa cell line was purchased from Cell Culture Facility at University of California, Berkeley, tested mycoplasma negative and authenticated with short tandem repeat analysis. The cells are maintained in DMEM supplemented with fetal bovine serum (10%) and penicillin/streptomycin (1%) in a humidified incubator at 37 ºC under 5% CO_2_. The 4-well plates and the *contact*Blot device are sterilized with 70% ethanol for at least 20 min in the tissue culture hood prior to use. Cells are detached from the tissue culture plate through trypsin treatment at ∼80% confluence, and resuspended in the fresh media to form a suspension of ∼1 million cells mL^-1^. The cell suspension is then filtered through a 35-μm membrane filter to eliminate cell clumps. After diluted down to ∼12% of the original concentration, the cell suspension is loaded onto the *contact*Blot devices, which are housed in 4-well plate chambers. After a ∼10-min cell settling period, the gels are washed gently with warm PBS to remove free cells that are not settled in microwells. Fresh media (∼6 mL) is added onto the device before the cell-loaded devices are placed in the CO_2_ incubator. The cells are cultured overnight on the device to recover from trypsin-release-induced stress and to form an adherent stance in each microwell. The duration of the incubation period is determined by the cell doubling time and the recovery rate of the adhesive bonds of the cells. For HeLa cells, the incubation duration is between 4 h (bond recovery time) and 24 h (doubling time). For the control experiments, cells from a suspension of the same density are cultured on a *contact*Blot device that is not functionalized with FN, and the same cell culture protocol is followed.

### Design of the *contact*Blot Assay

To assess the cell-contact state in culture, we perform a bright-field microscopy scan of the *contact*Blot device after HeLa cells are settled into the microwells and incubated for 1.5 h in the CO_2_ incubator. The brief incubation after cell settling allows weak bonds to form between each cell and the FN-patterned microwell, so that cells are adherent and immobilized during the scanning period. Using the ScanSlide function of the wide-field microscope, we scan an entire *contact*Blot device (25 × 75 mm) in 20 min at 10× magnification, thus providing sufficient resolution to examine the cell-contact state with a minimum perturbation on cell signaling. We then replace the cell-laden *contact*Blot device into the incubator for short-term culture of the cells in the FN-decorated microwells of the *contact*Blot device. After overnight culture (typically 8-12 h), HeLa cells form bonds with the microwell surfaces, and are ready for subsequent endpoint single-cell or contact-cell microwell-based western blot (μWB) experiments.^[32]^ Extracellular stimulation can be implemented at this stage, as described in our previous report.^[30]^ After immunoprobing, the abundance of the target proteins is measured from the fluorescence of the fluorescently-labeled antibody probes. Cell-contact state is inferred from the bright-field scan of the *contact*Blot device, with each cell’s state then indexed to the subsequent end-point μWB measurements to yield a multi-modal analysis of cell signaling in the context of cell-contact state.

### Application of the Osmotic Stress Stimulus

Isosmotic (300 mOsm) and hyperosmotic (500 mOsm) solutions are prepared by mixing 300 mOsm and 900 mOsm sucrose solution with cell culture media or PBS buffer. For *contact*Blot, HeLa cells are cultured overnight on the *contact*Blot device in the incubator before the implementation of the osmotic stress. Then the old medium is quickly removed from the device and replaced with an equal volume of osmotic solution. The cell-laden devices immersed in the osmotic solution are incubated in the CO_2_ incubator for 60 min to induce the osmotic responses. Upon the induction of the osmotic stress, the cells are analyzed for the abundance of the phosphorylated proteins through μWB as described above. To maintain a consistent osmotic condition, for every resuspension step, the cells are resuspended in the osmotic solution (PBS adjusted with the sucrose solution of the corresponding osmolarity). To mitigate potential phosphatase activity on the phosphorylated protein, a cocktail solution of phosphatase inhibitor is added to the lysis/electrophoresis buffer. Details for slab gel western blot of bulk cell suspensions can be found in Supporting Information.

## RESULTS AND DISCUSSION

### *Contact*Blot: A Multi-Mode Microfluidic Assay for Determining Phosphorylation Level of Isolated *versus* In-Contact Cells

To correlate heterogeneous cell behaviors with intracellular signaling response, we measure MAPK signaling in single HeLa cells in the context of neighboring cells. In this study, we extend our *in situ* single-cell western blot (*in situ* scWB) previously developed to measure the dynamic phosphorylation of MAPKs in stress-induced single-adherent cells.^[30]^ While microwells are similarly used to isolate cells in both studies, this study uses the microwells to facilitate cell-cell contact and the development of any associated intracellular signaling response during a short-term, on-chip cell culture period. The μWB device comprises ∼2000 microwells stippled in a thin-layer polyacrylamide gel mounted on a microscope slide (Figure 1A). Cells are gravity settled into the microwells (50 μm in diameter, ∼40 μm deep). To support short-term, in-well cell culture, each microwell is functionalized with the extracellular matrix protein fibronectin (Figure 2). In addition to forming the walls of the microwell features, during the end-point μWB, the polyacrylamide gel toggles between a protein-sieving matrix during protein electrophoresis to being a protein-capture scaffold (blotting membrane) upon brief exposure of the chip to UV light that activates benzophenone methacrylamide in the polymer network. Immobilized protein peaks are probed in-gel using primary and secondary fluorescently labeled antibody probes.^[32]^ To avoid disruptive dissociation of cell clusters, we integrate cell culture and western blotting on the same device for hundreds of individual cells, and hence we interrogate adherent cells in culture with single-cell resolution.

**Figure 1.**
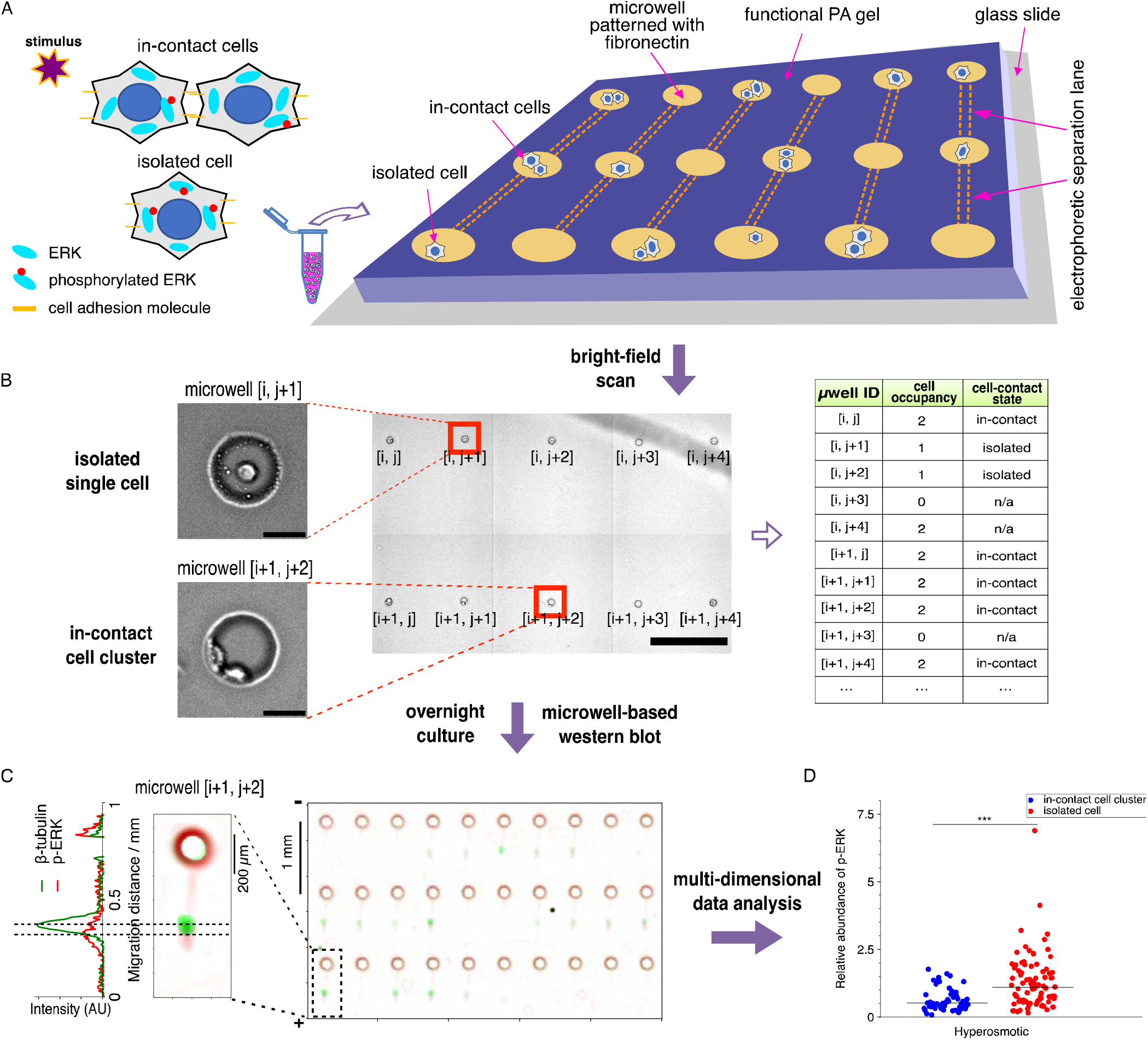
The multi-modal *contact*Blot assay prepends whole-cell bright*-*field imaging to determine cell contact state (isolated *versus* in-contact) with endpoint single-cell or cell-cluster western blotting (μWB) to measure ERK signaling. A microwell array format underpins concurrent on-chip culture and subsequent analysis of hundreds of isolated and in-contact cells. (A) To study ERK signaling in the context of cell-contact state, epithelial cells are loaded into microwells of a *contact*Blot device for on-chip cell culture. The microwell array is composed of ∼2000 microwells stippled in a polyacrylamide (PA) hydrogel layer. The PA gel first acts as the microwell walls and, subsequently, as a μWB protein electrophoresis and blotting gel. Cells are seated in each microwell, the floor of which is functionalized with fibronectin to support spreading of adherent epithelial cells. Microwells are 40 μm deep and 50 μm in diameter to accommodate both individual and clustered HeLa cells. (B) The cell-contact state is determined by bright*-*field scan (∼20 min) of the epithelial cell(s) accommodated in each microwell. After imaging, cells in the *contact*Blot are cultured on chip for 12 h before endpoint μWB of phosphorylated ERK (p-ERK) under an osmotic stress condition. Bright*-*field imaging of contact state and fluorescence imaging of the endpoint μWB are indexed to facilitate mapping back to the originating cell, thus reporting same-cell multi-modal profiles. Scale bars: left, 30 μm; middle, 500 μm. (C) Cell ERK signaling state is inferred from the level of p-ERK detected.^[30]^ (D) Comparative analysis of ERK signaling for isolated *versus* in-contact cells is conducted for each cell-occupied microwell by combining the imaging data with the *in situ* μWB results using the indexing framework.

**Figure 2.**
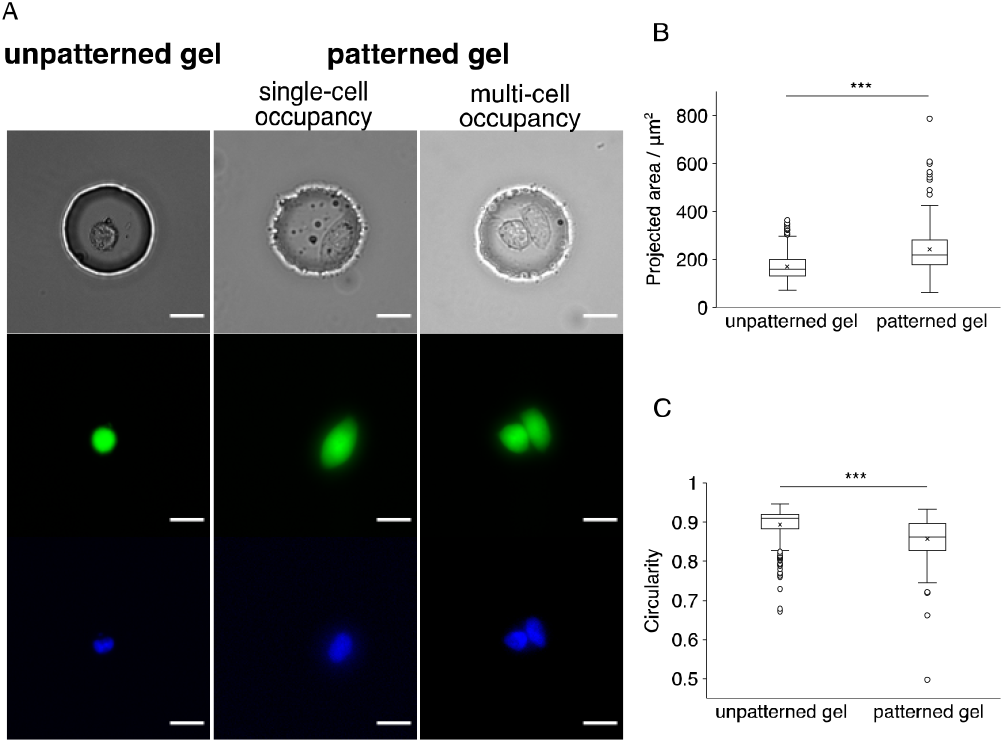
HeLa cells adhere and spread in microwells patterned with the extracellular matrix protein fibronectin (FN), but do not exhibit spreading in microwells that are not decorated with FN. (A) Bright*-*field and false-color fluorescence micrographs of HeLa cells cultured in microwells on a *contact*Blot device. Microwell dimensions: 50 μm in diameter, 40 μm in depth. Scale bars: 20 μm. FN concentration in the patterned gel: 10 μg ml^-1^. Green: calcein AM live stain. Blue: Hoechst33342 nucleic acid stain. HeLa cells were incubated for 12 h before imaging under the fluorescence microscope. (B) The projected area and (C) circularity of each cell or cell cluster on the unpatterned and patterned *contact*Blot devices. Circularity = 4π × (area/perimeter^2^). Statistical analysis is performed with the Mann-Whitney *U* test. \*\*\**P* < 0.001. Over 100 cells (305 cells for unpatterned gel, 206 cells for patterned gel) are analyzed for the experiment.

To perform comparative analysis of MAPK signaling on single isolated *versus* in-contact cells, the *contact*Blot assay comprises four steps, with details provided in the Supporting Information: (1) Gravity settle a cell suspension on the face of the open microfluidic device using an optimized cell-suspension density, so that the single-cell and two-cell microwell occupancies are dominant and sufficient for statistical analysis. (2) During a brief incubation period (∼1.5 h), adherent cells attach to the microwell walls through weak bonds with the fibronectin-decorated microwells. The cell-contact state of cells in each microwell is measured via bright-field microscopy inspection of the entire device (Figure 1B). This slidescan step can be repeated, acquiring state across multiple time points. The perturbation from the bright-field scan (∼20 min) can be recovered from an overnight incubation (Figure S1). (3) Application of osmotic stress for 60-min by buffer exchange, using an isosmotic (300 mOsm) or a hyperosmotic (500 mOsm) condition. (4) Completion of the μWB step, with indexing of the end-point μWB result for each cell-occupied microwell to the image-based cell-contact state (Figure 1C). In endpoint μWB quantitation, β-tubulin signal intensity was used for normalization to rule out the cell-size effect from protein-abundance analysis. In multi-cell occupancy, β-tubulin normalization also generates a cell-number-average result. Depending on cell-occupancy, the μWB reports targeted protein information from a single cell (single-cell occupancy) or an average of a cell cluster (multi-cell occupancy). Same-cell indexing allows data analysis to differentiate between signaling responses in the isolated cells *versus* the in-contact cells (Figure 1D). As such, we assess same-cell MAPK signaling and contact state simultaneously from the *contact*Blot for individual cells or cell pairs, depending on contact state.

### HeLa Cells Exhibit ERK Activation in Response to Osmotic Stress Regardless of Cell-Contact States

To validate the *contact*Blot assay, we examine ERK activation in response to hyperosmotic stress for isolated and in-contact cells. In HeLa cells, we have previously observed marked activation of the typical MAPKs (ERK, p38) induced by the hyperosmotic stress in single-cell and bulk experiments.^[30]^ Using the *contact*Blot assay, we measure both isolated cells and in-contact cells and – for both phenotypes – observe a significant increase in the level of the phosphorylated-ERK (p-ERK; normalized to β-tubulin) upon application of the hyperosmotic stress (Figure 3; Mann-Whitney *U* test, *n* > 100 blots, *P*_isolated_ = 2.139 × 10^−49^, *P*_in-contact_ = 1.065 × 10^−11^). In the isolated-cell group, the hyperosmotic-stress-induced increase of the median p-ERK is 6.4×, in agreement with current understanding.^[30, 33, 34]^ For the in-contact cell group under hyperosmotic stress, we observe a comparable increase in median p-ERK (6.1×). These observations suggest hyperosmotic stress induces similar ERK-activation regardless of the cell-contact state.

**Figure 3.**
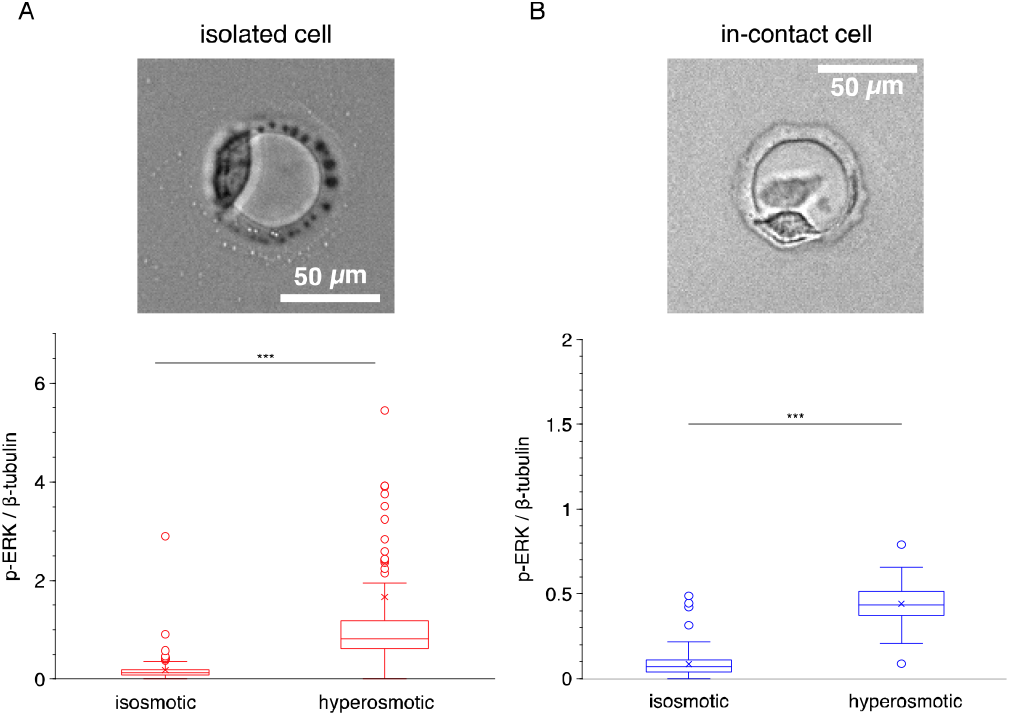
*Contact*Blot reports ERK activation levels of HeLa cells in response to hyperosmotic stress that are similar for both isolated and in-contact cells. (A-B) Osmotic response of single isolated (A) and in-contact (B) HeLa cells via ERK signaling. Top, repres*e*ntative bright-field micrographs (10× objective) of HeLa cells cultured in the microwell. Bottom, box plots showing the β-tubulin-normalized p-ERK levels from μWB measurements. Over 100 μWB assays are analyzed for each μWB experiment. Upon a 60-min application of the hyperosmotic stress of 500 mOsm, a significant increase of p-ERK is detected regardless of the cell-contact states, with the increase fold of 6.4× and 6.1× in the median p-ERK level for isolated and in-contact cells, respectively. Statistical analysis is performed with the Mann-Whitney *U* test. \*\*\**P* < 0.001. Scale bars: 50 μm.

To validate the μWB assessment, we perform bulk western blotting of cell groups, where we tune the cell confluency to represent the cell-contact state (Figure 4). In the population-averaged slab-gel western-blot measurements, we observe a significant increase in the p-ERK level (normalized to β-tubulin) induced by the hyperosmotic stress condition in both low-confluence (2.5 × 10^4^ cells) and high-confluence (20 × 10^4^ cells) groups (one-tailed Student’s *t*-test, *n* > 3, *P*_low-confluence_ = 4.503 × 10^−4^, *P*_high-confluence_ = 2.557 × 10^−6^). In sum, comparable ERK-activation stress-response levels are observed in the population-wide groups, regardless of cell-contact state.

**Figure 4.**
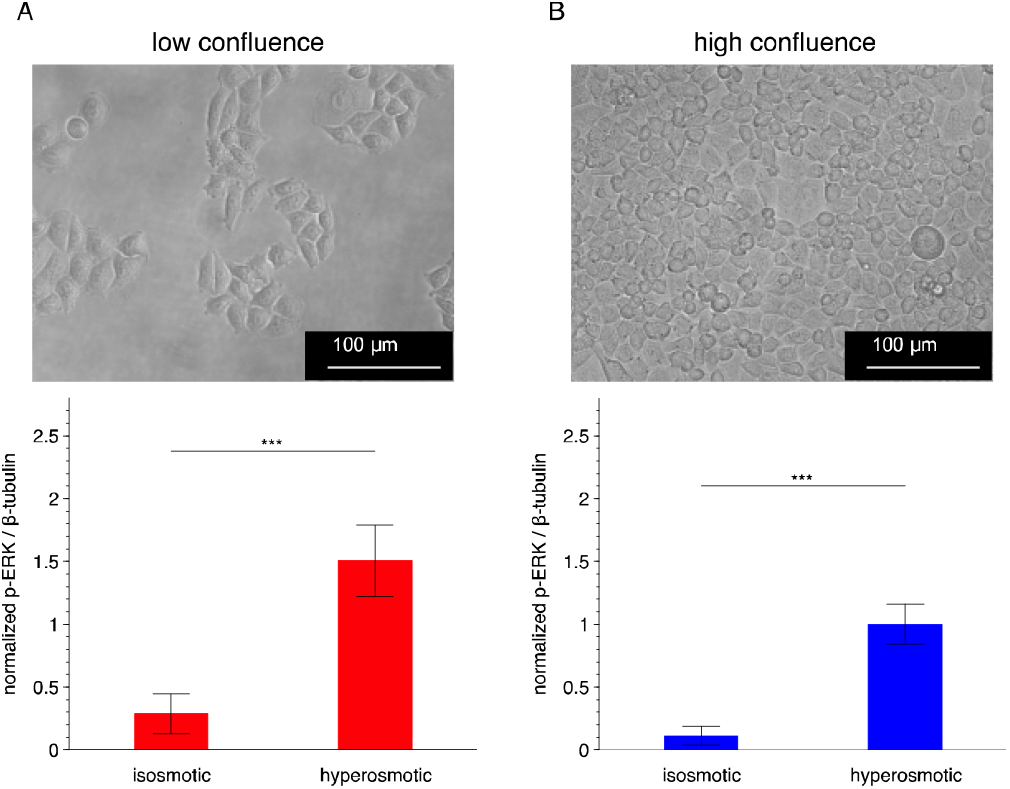
ERK activation levels of HeLa cells in response to hyperosmotic stress *in bulk*, similar for both isolated and in-contact cells. Osmotic response of low-confluence (*A*) and high-confluence (*B*) HeLa cells in ERK signaling. Top, representative bright*-*field micrographs (20× objective) of HeLa cells cultured in the bulk experiments. The seeding cell numbers are 2.5 × 10^4^ and 20 × 10^4^ for low- and high-confluence experiments, respectively. Bottom, bar plots showing the β-tubulin-normalized p-ERK levels from bulk measurements. Data are presented as means ± SDs (*n* > 3). Upon a 60-min application of the hyperosmotic stress condition, a significant increase of p-ERK is detected regardless of the cell-confluence level. Statistical analysis is performed with the one-tailed Student’s *t*-test. \*\*\**P* < 0.001. Scale bars: 100 μm.

The validation study suggests that *contact*Blot reports cellular-resolution ERK phosphorylation under hyperosmotic stress, as is consistent with previous reports. Furthermore, *contact*Blot shows comparable ERK-activation levels between the isolated *versus* in-contact cells in response to hyperosmotic stress.

### HeLa Cells Exhibit Attenuated ERK Signaling in the In-Contact Cell Group

Having validated the *contact*Blot assay on hyperosmotic-stress-induced ERK activation, we next seek to understand the role of cell-cell contact on signal transduction. Cell-cell contacts have been largely reported to inhibit the activity of epithelial cells in bulk.^[1—8]^ Because of the clustering tendency of adherent cells, analyzing cells in bulk provides limited insight. Using precision microfluidic analytical tools, we aim to examine the ERK signaling with unambiguously isolated and in-contact cells to gain a more precise understanding of the role of cell-contact state.

We first examine the p-ERK levels under the isosmotic condition for both isolated cells and in-contact cells (Figure 5A). We make two key observations: (1) the median p-ERK level in the in-contact cell group is significantly lower than in the isolated-cell group (Mann-Whitney *U* test; *n* > 100 cells, *P* = 7.695 × 10^−11^ for replicate 1, *n* > 50 cells, *P* = 1.008 × 10^−9^ for replicate 2), and (2) the median p-ERK level in the in-contact cell group is ∼50% of that same value in the isolated-cell group (0.56× for replicate 1, 0.42× for replicate 2).

**Figure 5.**
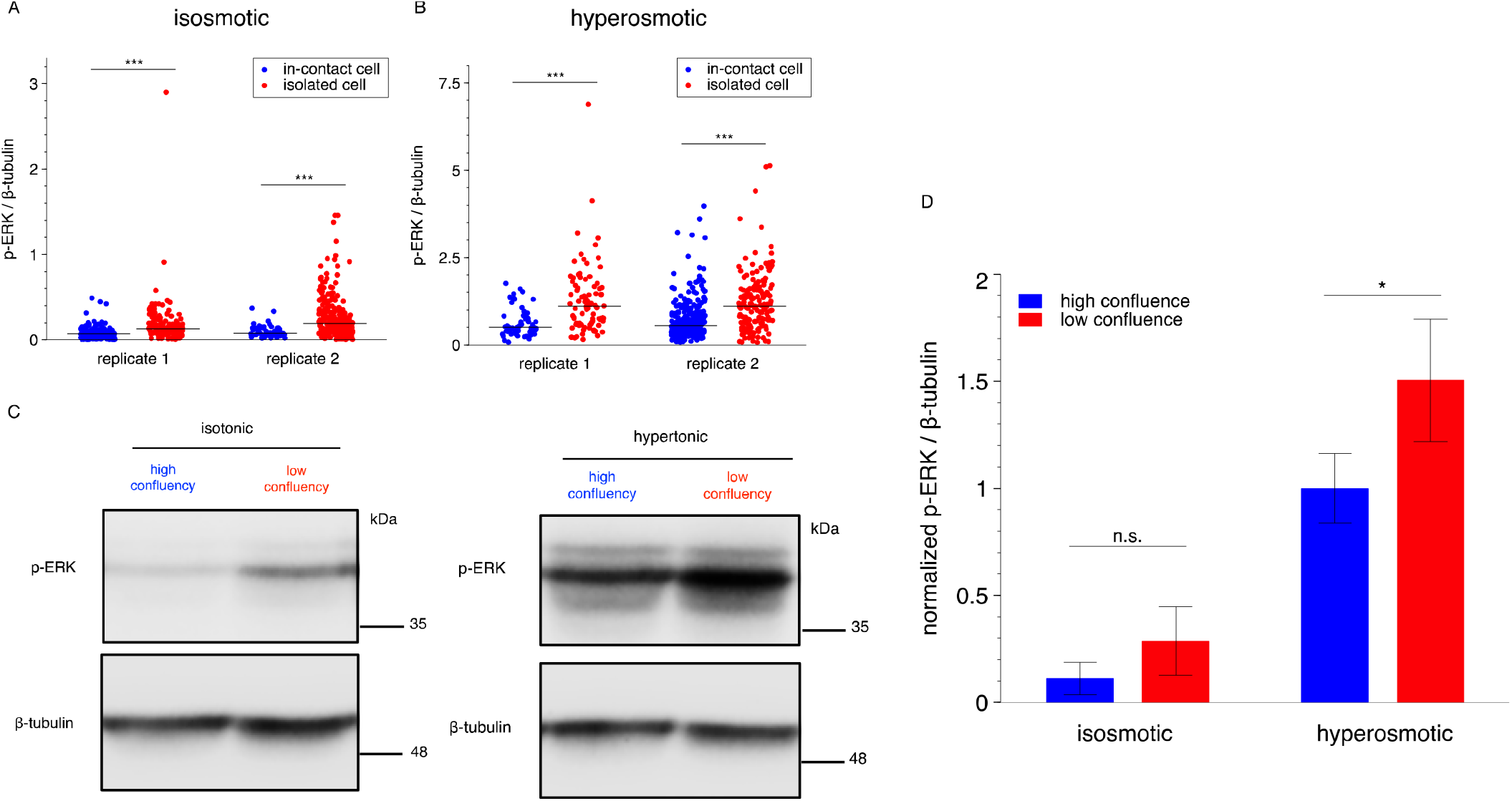
*contact*Blot reports differential ERK activation between the isolated and in-contact HeLa cells. (A-B) HeLa cells exhibit a significantly attenuated ERK activation level in in-contact *versus* isolated single cells. The contact-attenuation in ERK activation is observed under both the isosmotic (A) and hyperosmotic (B) conditions. Solid lines across the data sets denote the median level. Statistical analysis is performed with the Mann-Whitney *U* test. \*\*\**P* < 0.001. Over 100 μWBs are analyzed for each μWB experiment. (C) Bulk western blotting of a population of HeLa cells confirms the attenuated activation of ERK in high-confluence cells (full-size gels in Supporting Information). The seeding cell numbers are 2.5 × 10^4^ and 20 × 10^4^ for low- and high-confluence experiments, respectively. β-tubulin is used as a loading control. (D) Quantitative analysis of ERK differential activation in low- and high-confluence cells from the population-level western blotting experiments. The difference in the ERK activation levels between the cells in high- and low-confluence is not significant under the isosmotic condition and marginally significant under the hyperosmotic condition, yet less pronounced than observed with *contact*Blot. Data are presented as means ± SDs (*n* > 3). Statistical analysis is the one-tailed Student’s *t*-test. \**P* < 0.05; n.s., *P* > 0.05.

phorylation levels at the end of the 60-min hyperosmotic shock. By comparing the p-ERK levels between the isolated-cells and in-contact cells, analysis can shed light on the differential phosphorylation levels of ERK as related to cell-contact state, after the hyperosmotic treatment (Figure 5B). We observe that the median p-ERK level in the in-contact cell group is significantly lower than the value measured in the isolated-cell group (Mann-Whitney *U* test; *n* > 50 cells, *P* = 1.496 × 10^−6^ for replicate 1, *n* > 150 cells, *P* = 4.055 × 10^−16^ for replicate 2). We further observe that the differences in the median p-ERK level between the isolated and the in-contact cell groups are 0.45× and 0.50× for replicate 1 and 2, respectively.

In contrast, in bulk assays (where no physical confinement is imposed) HeLa cells tend to grow in clusters even at a low seeding density (Figure 4A), thus presenting challenging ambiguity in analyses of isolated (no cell-cell contact) cells. Consequently, assessing the difference in p-ERK levels between isolated and in-contact cells is challenging using bulk cell culture approaches. For comparison, we analyze the bulk data in Figure 4, and observe a lower p-ERK level in the highly confluent cells (assuming mostly in-contact cells) compared to the Next, to understand the impact of cell-cell contact on the acute stress responses, we investigate ERK phoslow-confluence cells (Figure 5C and 5D), but the difference is less pronounced as compared to the degree of difference observed in *contact*Blot experiments (one-tailed Student’s *t*-test, *n* > 3 replicates; *P* = 0.015 for the hyperosmotic condition, *P* = 0.056 for the isosmotic condition). We attribute the subtle difference observed in the slab-gel western blot to i) the inclusion of a population of in-contact cells in even the low-confluence group and ii) insufficient detection sensitivity for lower abundance protein targets under the isosmotic conditions in the low-confluence group. Two confounding factors that can be readily addressed with enhanced precision in *contact*Blot in cell handling and analysis.

Taken together, we observe an attenuated level of p-ERK in the in-contact cell group under both hyperosmotic and isosmotic conditions. The differential levels of p-ERK between the isolated and in-contact cells are distinguishable using the precision *contact*Blot because: i) cell confinement in microwells enforces an isolated cell state as compared to standard plate cell culture, ii) *in-situ* analyses during cell culture reduce perturbations from cell-sample preparation and handling, and iii) relatively weak signals from the endpoint μWB of single, isolated cells under the isosmotic condition are detectable using the photomultiplier tube-based fluorescence detection.

While this study focuses on ERK activation given relevance in regulating cell proliferation, the interplay of MAPKs in coordinating cell responses to environmental stimuli suggests potential differential activation for other kinases in the MAPK family, such as p38 and JNK. Attenuated levels of p-p38 and p-JNK were observed in fibroblasts in bulk experiments.^[35]^ Our bulk data support this hypothesis with HeLa cells (Figure 6). More broadly, enzymes that participate in the regulation of MAPKs, such as tumor necrosis factor receptor 1 (TNFR1), MAP kinase kinase (MKK), and MAP kinase phosphatase (MKP), may also be candidates expressing differential activation levels between isolated and in-contact cellular conditions.

**Figure 6.**
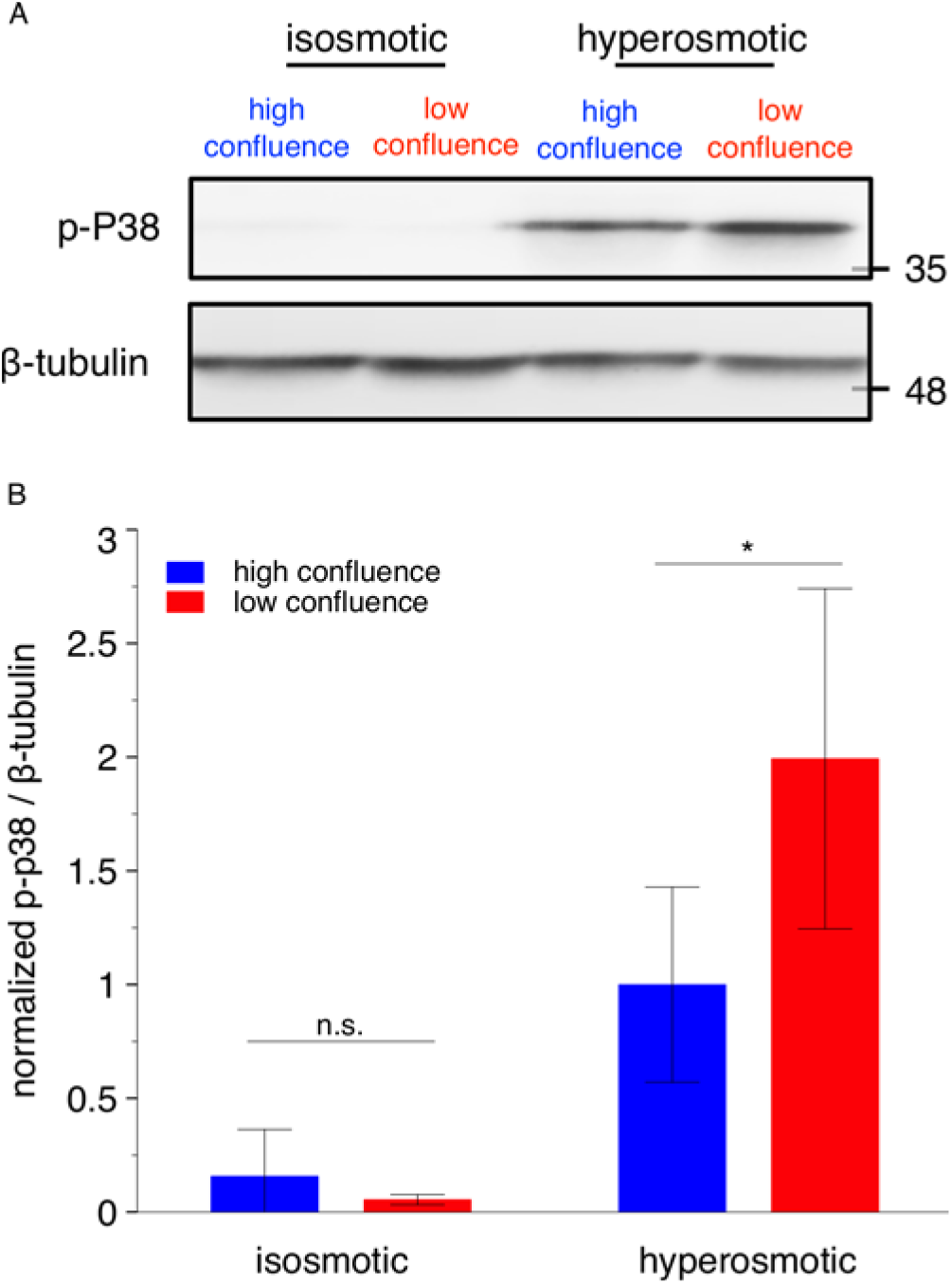
Differential activation of p38 in response to hyperosmotic stress between HeLa populations of different confluence levels was observed in bulk. (A) Western blotting of a HeLa cell suspension confirms the attenuated activation of p38 in high-confluence cells in response to the hyperosmotic stress. The seeding cell numbers are 2.5 × 10^4^ and 20 × 10^4^ for low- and high-confluence experiments, respectively. β-tubulin is used as a loading control. The isosmotic and hyperosmotic conditions are 60 min of 300 and 500 mOsm, respectively. The western blotting for isosmotic and hyperosmotic conditions is performed on the same membrane and under the same detection condition. (B) Quantitative analysis of the activation of p38 in low- and high-confluence cells from the population-level western blotting experiments. The difference of the p38 activation levels between the cells in high- and low-confluence is not significant at the isosmotic level, and slightly significant at the hyperosmotic level. Data are presented as means ± SDs (*n* > 3). Statistical analysis is performed with the one-tail Student’s *t*-test. \**P* < 0.05; n.s., *P* > 0.05.

The observed attenuation of p-ERK in the in-contact cells could very likely be a combined effect involving both the physical cue of cell-cell contact and the chemical cues emanating from proximal cells. In fact, the change of radical levels related to cellular metabolic activities can alter the local redox environment. The alteration in local redox environment correlates with contact-inhibited proliferation across cell lines.^[36, 37]^ Future studies could include finer classification of cell-cell interactions (i.e., consideration of the cell-cell distance in the isolated-cell group, profiling of secreted factors) to create a holistic understanding of the milieu created during cell-cell contact.

## CONCLUSION

We introduce, validate, and apply an imaging-facilitated *in situ* cellular-resolution western blot (*contact*Blot) assay that reports cell-contact information and protein signaling activity for an array of hundreds of individual cells. This multi modal analysis tool provides comparative targeted protein analyses of cancer cells, both in isolation and in contact with neighboring cells. Such profiling may clarify differential protein levels in early and late phases of tumor development, hence facilitating analysis of new cancer-drug targets related to cell-cell contact. Differential stress responses of isolated and in-contact cells may delineate therapeutic susceptibility of diseased cells at different stages in disease progression, thereby informing treatment decisions. Furthermore, by including various fluorescent probes in imaging, the *contact*Blot assay indexes the μWB-derived targeted protein signatures of individual cells to a broader range of physical and biochemical conditions of the same cells, including intracellular temperature,^[38, 39]^ membrane potentials,^[40]^ O_2_ levels,^[41]^ pH levels,^[42]^ and cell-cycle phases.^[43]^

## ASSOCIATED CONTENT

### Supporting Information

Additional experimental details, including reagents, device fabrication, assay protocols, statistics, supplementary figures, tables, and discussions (PDF).

### Author Contributions

The manuscript was written through contributions of all authors.

## ACKNOWLEDGMENT

This work was supported by the US National Institutes of Health (R01CA203018 and R33CA225296 to A. E. H.), the Chan Zuckerberg Biohub (to A. E. H.), and JSPS KAKENHI (Grant Number JP15KK0297, JP18H02569, and JP22H02761 to I. N.). We thank the QB3 Biomolecular Nanotechnology Center in Stanley Hall and the CRL Molecular Imaging Center (supported by the Helen Wills Neuroscience Institute) at the University of California, Berkeley. We appreciate helpful discussions with Herr Lab members throughout this project. Henrietta Lacks, and the HeLa cell line that was established from her tumor cells without her knowledge or consent in 1951, have made significant contributions to scientific progress and advances in human health. We are grateful to Henrietta Lacks, now deceased, and to her surviving family members for their contributions to biomedical research. The content is solely the responsibility of the authors and does not represent the official views of the National Institutes of Health.

