## Supplementary Information for "*Contact*Blot: Microfluidic Control and Measurement of Cell-Cell Contact State to Assess Contact-Inhibited ERK Signaling"

**Abstract:** Extracellular signal-regulated kinase (ERK) signaling is essential to regulated cell behaviors, including cell proliferation, differentiation, and apoptosis. The influence of cell-cell contacts on ERK signaling is central to epithelial cells, yet few studies have sought to understand the same in cancer cells, particularly with single-cell resolution. To acquire same-cell measurements of both phenotypic (cell-contact state) and targeted-protein profile (ERK phosphorylation), we prepend high-content, whole-cell imaging prior to endpoint cellular-resolution western blot analyses for each of hundreds of individual HeLa cancer cells cultured on that same chip, which we call contactBlot. By indexing the phosphorylation level of ERK in each cell or cell-cluster to the imaged cell-contact state, we compare ERK signaling between isolated and in-contact cells. We observe attenuated (~2×) ERK signaling in HeLa cells which are in-contact versus isolated. Attenuation is sustained when the HeLa cells are challenged with hyperosmotic stress. Our findings show the impact of cell-cell contacts on ERK activation with isolated and in-contact cells, while introducing a multi omics tool for control and scrutiny of cell-cell interactions.

[a]	Dr. Y. Zhang, Prof. A. E. Herr
Department of Bioengineering
University of California, Berkeley
Berkeley, CA, 94720, USA


[b]	Dr. I. Naguro, H. Ryuno
Graduate School of Pharmaceutical Sciences
The University of Tokyo
Tokyo, Japan

[c]	Dr. I. Naguro,
Faculty of Pharmacy
Juntendo University
Chiba, Japan

### Experimental Procedures

*Reagents:*

Ammonium persulfate (APS; A3678), N,N,N’,N’-tetramethylethylenediamine (TEMED; T9281), acrylamide/bisacrylamide solution (30% 37.5:1; A3699), Tris-HCl (pH 6.8; BBT-403; Boston Bioproducts), Tris-HCl (pH 8.8; T1588; Teknova), microscope glass slide (25 mm x 75 mm x 1mm; 48300-048; VMR), fibronectin (bovine plasma powder; F4759), N-[3-[(3-benzoylphenyl)formamido]propyl]methacrylamide (BPMAC, PharmAgra), Deionized water (ddH2O; 18.2 mΩ; Millipore), dichlorodimethylsilane (440272), 3-(trimethoxysilyl)propylmethacrylate (440159), sodium dodecyl sulfate (SDS; L3771), sodium deoxycholate (D6750), Triton X-100 (X100), 10x Tris-glycine (25 mM, pH 8.3; 161-0734; Bio-Rad), phosphatase inhibitor cocktail (PI78442; FisherScientific), RIPA buffer (89900, ThermoFisher), bromophenol blue (B0126), glycerol (G2025), dithiothreitol (D0632), sucrose (S0389), bovine serum albumin (A7030), Tris-buffered saline with Tween 20 (TBST, 10×; 9997S; Cell Signaling Technology), primary antibodies for β-tubulin (rabbit; ab6046; Abcam), for phosphorylated extracellular signal-regulated kinase (p-ERK) (rabbit; 4370S; Cell Signaling Technology), and for phosphorylated p38 (p-p38) (rabbit; 4511S; Cell Signaling Technology), anti-rabbit secondary antibody (Donkey, AlexaFluor 555; A31572; Life Technologies), 2-mercaptoethanol (M3148), DMEM (DMEM(1×) + GlutaMAX-1 supplement; 10566-016; Gibco), fetal bovine serum (FBS; sterile; 100106; Gemini), penicillin/streptomycin (Pen Strep; 15140-122; Gibco), MEM non-essential amino acids solution (MEM NEAA; 100×; 11140050; Life Technologies), phosphate buffered saline (PBS; sterile, pH 7.4; 10010023; Gibco), Trypsin-EDTA (0.05%; 25300-054; Gibco), LIVE/DEAD® Viability/Cytotoxicity Assay Kit (L3224; Life Technologies), Hoechst 33342 (H3570; Life Technologies), cell strainer (sterile; 352235; Corning), silicon wafer (100-mm diameter; C04009; WaferPro), SU-8 photoresist (3050; Microchem), SU-8 developer (Y020100; Microchem), mylar mask with microwell features (CAD/Art Services). All the chemicals and reagents that are not specified with the manufacturer information were purchased from Sigma.

*Microfabrication:*

Fabrication of SU-8 mold: To cast the polyacrylamide (PA) gel for the *contact*Blot device, we fabricate the SU-8 mold through a standard photolithography protocol.^[1]^ The SU-8 mold is designed to create arrays of micro-posts (50 𝜇m in diameter and 40 𝜇m in height) for molding of microwells dimensioned to allow for HeLa cell spreading in the microwell during the overnight culture. Microwell pitch (~500 𝜇m) is designed to reduce chemical-signal crosstalk between neighboring cell-laden microwells, while maintaining a substantial number of microwells on each glass microscope slide.

Cell spreading assessment: The evaluation of cell spreading is performed by examining fluorescence micrographs of calcein AM-stained cells. The staining protocol is similar to a viability validation assay, but the staining solution includes only calcein AM. Specifically, the cell-laden device is incubated in calcein AM solution (10 𝜇M in PBS) in the dark at room temperature for ~20 min. At the end of the staining, the device is washed off the staining solution with warm PBS for fluorescence imaging. To ensure a precise calculation of the cell contours, fluorescence micrographs are obtained at a high magnification (20×). Each micrograph of the cell is quantitated for the projected area and circularity (circularity = 4𝜋 x (area/perimeter^2^)).

*Recovery culture after ScanSlide:*

When measuring ERK signaling, a recovery cell-culture period is needed to restore the baseline (under the isosmotic condition) ERK phosphorylation levels after the brightfield scan of the device, which is housing living HeLa cells during the scan period. We observe that a 2-h incubation following the scan yields phosphorylated ERK (p-ERK) under the isosmotic condition remains at a high level comparable to the p-ERK under the hyperosmotic condition (Figure S1A). Whereas after overnight incubation following the scan, p-ERK under the isosmotic condition decreases to a normal level (Figure S1B).^[2-4]^ We attribute the post-scan increase in p-ERK under the isosmotic condition to the perturbations during the scanning process, including the changes in temperature and CO_2_ level, and potentially fluid-motion associated stress induced by the frequent movement of the microscope stage. A microscope equipped with an environmental chamber would reduce the time required for the recovery cell-culture by maintaining the temperature and CO_2_ level during slide scanning.

*Osmotic stress application:*

For slab gel western blot of bulk cell suspensions, cells are cultured in 24-well plates (500 μL medium in each well) for 1 to 2 days before osmotic-stress induction. Then 250-μL osmotic solution (300 or 900 mOsm) is added to the cell culture in each well. Following an incubation of a defined time, the medium in each well is aspirated and the ice-cold lysis buffer (100 μL, RIPA buffer + Protease/Phosphatase inhibitor cocktail) is immediately added to the well. After a 20-min 4 ºC incubation on the shaker, the cell lysates are transferred to Eppendorf tubes for centrifugation (13500 rpm, 5 min, 4 ºC). Upon centrifugation, 50 𝜇l of supernatant in each tube is transferred to a new tube containing 50 𝜇l lab-made 2× SDS sample buffer (100 mM Tris-HCl pH8.8, 10%w/v bromophenol blue, 36%v/v glycerol, 4%w/v sodium dodecyl sulfate, 10 mM dithiothreitol). The samples are denatured at 98 ºC for 3 min and subjected to western blot analysis.

*Microscopy:*

Cell spreading and cell contact state are assessed via widefield imaging. An Olympus IX71 inverted fluorescence microscope (equipped with ASI motorized stage, X-cite mercury lamp light source (Lumen Dynamics), and standard FITC and Cy5 filter cube (10× and 20× objectives)) is used for imaging. The bright-field and fluorescence micrographs are obtained using an iXon+ EMCCD camera (Andor Technology Ltd.) controlled by a MetaMorph software (Molecular Devices) with 10 ms exposure time. The fluorescence images of the protein blots in 𝜇WB experiments are collected by scanning the *contact*Blot device with a fluorescence microarray scanner (Genepix 4300A, Molecular Devices).

*Data analysis:*

Data analysis for 𝜇WB experiments is performed with Matlab (MathWorks) using an in-house analysis script.^[1]^ Briefly, background-subtracted signals of the proteins are integrated within the defined boundaries for AUC calculation. As a quality control, the protein peaks with SNR ≥ 3 are analyzed. All the other data analysis is performed with Fiji (2.3.0/1.53f, https://imagej.net/software/fiji/).

*Statistical analysis:*

Mann-Whitney *U* test is used to evaluate the difference between the groups that do not follow the normal distribution, including the comparisons of the normalized phosphorylated protein abundance between the populations of iso- and hyper-osmotic conditions, the comparisons between the isolated and in-contact cells, and the comparisons of the spreading levels between the cells cultured on FN-patterned and unpatterned *contact*Blot devices. One-tailed student’s *t* test is used to compare the phosphorylation levels of ERK and p38 between the cells under different osmotic conditions or under different contact states in bulk experiments.

### Results and Discussion

The observation of differential ERK signaling between the isolated and in-contact cells under isosmotic conditions is in alignment with the growth-promoting function of ERKs. As a ubiquitous MAPK protein responding to growth factors (e.g., EGF), activated ERKs promote cell-cycle entry by regulating the expression of Cyclin D1.^[5, 6]^ Hence, the inhibited growth and division in in-contact cells would involve a shift of the ERK activation state from the state in isolated cells. In fact, lower p-ERK levels in high-density cells are common in epithelial cell lines,^[7-9]^ consistent with our observations. Interestingly, under hyperosmotic stress we observe differential ERK signaling between isolated- and in-contact cells, and the difference in the median level of p-ERK is comparable to the difference at the isosmotic level. Combining with the comparable hyperosmotic-stress-induced ERK activation levels between the isolated- and in-contact cells, our study suggests that 1) under the hyperosmotic condition, the observed differential p-ERK levels between the isolated- and in-contact cells arise from the attenuated p-ERK level in the in-contact cells occurring in culture and 2) an acute osmotic shock generates comparable fold changes in ERK response in isolated- and in-contact cells.

Contact-inhibited cell behavior has been extensively studied in normal epithelial cells (e.g., fibroblasts, MCF-10A cells, Madin-Darby canine kidney (MDCK) cells).^[5, 7, 9]^ Cancer cells are typically considered to lack contact-inhibition even in high-confluence populations.^[10]^ Reports have suggested that the loss of contact-inhibition features may be a hallmark of cancer.^[7, 11]^ However, inhibited growth of HeLa cells was noticed in an early microscopic study of cellular contact behaviors, where piling or strong overlapping of cells was not detected.^[12, 13]^ In addition to HeLa cells, the typical breast cancer cell line MCF-7 has been reported to exhibit contact-inhibited proliferation.^[14]^ By identifying the existence of contact-attenuated p-ERK in HeLa cells, our findings, along with other contact-inhibition studies on cancer cells, suggest that contact-inhibited activity exists in certain cancer cell lines, and loss of contact-inhibition may not be a generic hallmark for cancer cells, at least not in some *in vitro* cultured cancer cell lines. Understanding the regulation of cancer cell activities in the context of cell-cell contacts would facilitate molecular studies of tumor progression.

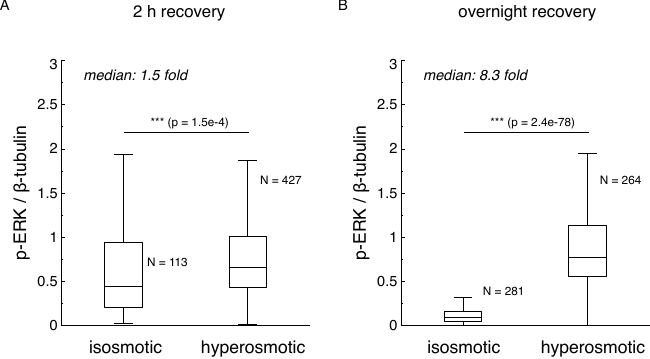
**Figure S1.** The perturbation on the phosphorylation measurement of HeLa cells from the bright-field scan can be recovered after an overnight recovery culture. (A) The phosphorylated level of the extracellular signal-regulated kinase (p-ERK) under the isosmotic condition is high when examined after 2-h incubation from the bright-field scan of the *contact*Blot device. (B) The p-ERK level under the isosmotic condition decreases to a normal level compared to our previous study^[2]^ and the reports in bulk measurements,^[3, 4]^ when examined after 12-h incubation from the brightfield scan. Statistical analysis is performed with the Mann-Whitney *U* test. ****P* < 0.001. The isosmotic and hyperosmotic conditions are 60 min of 300 and 500 mOsm, respectively.

**Figure S2.** Uncropped slab-gel western blot images for Figure 5C.

### Author Contributions

Y. Z. and A. E. H. designed research; Y. Z., I. N., and H. R. performed research; Y. Z. and I. N. analyzed data; and Y. Z. and A. E. H. wrote the paper.
